# The Dynein-Dynactin complex influences amnioserosa cell morphodynamics and adhesion during Drosophila dorsal closure

**DOI:** 10.1101/2025.10.21.683620

**Authors:** Deepanshu Sharma, Maithreyi Narasimha

## Abstract

The contraction of the amnioserosa, which provides the major force for Drosophila dorsal closure, is powered by apical constriction. While much work has demonstrated the requirement for the spatiotemporal regulation of intracellular forces generated by the actomyosin cytoskeleton in patterning apical constriction, the role of the microtubule cytoskeleton remains poorly understood. We previously uncovered the necessity of apical microtubule meshwork reorganisation in patterning apical constriction dynamics, and demonstrated its reliance on EB1-dependent microtubule growth and Patronin-dependent non-centrosomal nucleation. Here we discover the requirement for Dynein-Dynactin complex functions in patterning cell morphodynamics in the amnioserosa. We show that perturbing this complex in multiple ways, including the use of a mutant that likely disrupts microtubule anchorage rather than transport, alters apical constriction morphodynamics and modulates adherens junction remodelling during amnioserosa cell delamination. The defects we observe are additionally sensitive to ECadherin levels and lead to compromised amnioserosa integrity and adverse dorsal closure outcomes. We suggest that the Dynein-Dynactin complex influences cell morphodynamics in the amnioserosa by enabling the regulation and spatiotemporal coordination of cell adhesion, the mechanistic basis for which remains to be discovered.

## Introduction

Drosophila dorsal closure serves as a tractable model for a class of morphogenetic movements termed ‘epithelial fusion’ that includes neural tube and palatal closure in vertebrates, and accomplishes the establishment of continuity between two epithelial flanks initially separated by an intervening tissue. Decades of pioneering work have informed our understanding of the tissue contributions and cell behaviours that enable timely and precise dorsal closure (Kiehart *et al*., 2000; Jacinto *et al.,* 2002; Hutson *et al*., 2003; Narasimha and Brown, 2004; Toyama *et al*., 2008; Solon *et al*., 2009; Das Gupta and Narasimha, 2019). Contraction of the intervening tissue, the amnioserosa, by apical constriction brings the dorsal epidermal flanks that are attached to its perimeter by cell-cell and cell-substrate adhesion closer to each other, where they form new contacts and, through it, establish epithelial continuity. In the epidermal flanks, transient reorganisation of dynamic microtubules in the leading edge epidermal cells facilitates ‘zipping’ at the canthi at the anterior and posterior ends of the eye-shaped amnioserosa. The assembly of a supracellular actin cable at the interfaces of the leading-edge cells in contact with the amnioserosa, although dispensible for force generation, ensures coordination by maintaining the right tissue geometry (Jankovics and Brunner, 2006; Ducuing and Vincent, 2016; Pasakarnis *et al*., 2016).

Apical constriction in the amnioserosa is spatiotemporally heterogeneous: it is pulsed and slow during early closure (Phase I) and unpulsed and rapid in its latter part (Phase II). In addition to this, cell delamination, the basal exit of single cells from the amnioserosa, is a seemingly stochastic and relatively rare cell behaviour accompanied by rapid apical constriction. Thus, the amnioserosa is an excellent in-vivo test bed in which to interrogate the molecular and physical basis of heterogeneities in cell behaviour, specifically the nature, regulation, and coordination of intracellular forces that power them. While much work has focused on the organisation of the actomyosin cytoskeleton in the contributing tissues that underlie heterogeneities in cell behaviour (Gorfinkiel *et al*., 2009, 2025; Solon *et al*., 2009; Blanchard *et al*., 2010; Meghana *et al*., 2011; Saravanan, *et al.,* 2013; Duque and Gorfinkiel, 2016), the regulation and requirement of the microtubule cytoskeleton remain relatively poorly understood.

We recently uncovered the requirement for the reorganisation of the apical microtubule meshwork in patterning apical constriction in the amnioserosa and demonstrated its reliance on EB1-dependent microtubule growth and patronin-dependent nucleation. Specifically, we showed that reorganisation is sensitive to the reduction of patronin: in a subset of patronin hypomorphic mutants, the apical microtubule meshwork appeared to be ‘anchored’ to the apical cortex, and mutant amnioserosa cells exhibited reduced constriction pulse amplitudes (Guru *et al*., 2022). This raised the possibility that microtubule anchorage to the cortex might be regulated spatiotemporally during apical constriction. In the nearest neighbours of delaminating amnioserosa cells, which we previously showed contribute forces for delamination, microtubules were observed to realign and polarise with their plus ends directed towards their interfaces with the delaminating cell (Meghana *et al*., 2011; Saravanan *et al.,* 2013). This suggested that microtubule anchorage, potentially through the plus end, might also contribute to the dynamics and mechanics of cell delamination.

Microtubule anchorage to the plasma membrane/ cortex can occur through its plus or minus ends. While Patronin and the Gamma tubulin ring complex (γTuRC) mediate minus-end-dependent anchorage (Akhmanova and Kapitein, 2022), NuMA/Mud has been shown to anchor microtubules through microtubule plus ends. Mud links the Dynein-Dynactin complex on microtubule plus ends with Pins (the Drosophila homolog of LGN) and G proteins at the apical membrane (Kiyomitsu and Boerner, 2021) to anchor astral microtubules to the cortex and mediate spindle attachment and positioning in dividing cells (Izumi *et al*., 2006). Mud is also enriched at the tricellular junctions of epithelial cells, where they enable cells to retain a molecular memory of their interphase cell shape after cell division (Bosveld *et al*., 2016). However, the role of Mud or Dynein-Dynactin complex-dependent microtubule anchorage in facilitating cell shape changes or modulating cell deformability in non-dividing cells remains poorly investigated.

The Dynein-Dynactin complex is a multi-subunit complex that is necessary for intracellular functions that include cargo transport, organelle positioning, microtubule organisation, and anchorage. Some perturbations that interfere with the organisation of the Dynactin complex can disrupt microtubule architecture without necessarily affecting transport, suggesting that the two functions can be uncoupled (Kim *et al*., 2007). Key structural components of the Dynactin complex include the actin-related protein 1 (Arp1), the homodimeric protein Dynamitin/p50, and the p150/Glued subunit. Recent studies have identified a neomorphic mutation in the Arp1 subunit of the complex that likely selectively affects microtubule anchorage and growth without affecting transport, and demonstrated its influence on the subcelluar enrichment of proteins and RNAs in the Drosophila oocyte (Nashchekin *et al.,* 2016). Dynamitin serves as a crucial scaffold in the Dynein-Dynactin complex by linking Arp1 (the core filament) and p150 (the microtubule binding subunit). Overexpression of Dynamitin functions as a dominant-negative inhibitor that results in dissociation of the complex (Echeverri *et al*., 1996; Melkonian *et al*., 2007).

Here, we combine genetics with high-resolution real-time confocal microscopy to examine the influence of the Dynactin complex on cell morphodynamic heterogeneities that accompany amnioserosa contraction during Drosophila dorsal closure. Using multiple ways of perturbing the complex, including a mutant that selectively affects microtubule anchorage, we uncover a role for the complex in modulating cell deformability during apical constriction and cell adhesion remodelling during cell delamination.

## Results

We chose to examine the morphogenetic roles of the Dynein-Dynactin complex in the amnioserosa which is post-mitotic and performs a purely mechanical function of providing forces during dorsal closure through patterned apical constriction. To enable the perturbation of the microtubule anchorage functions of the complex rather than its transport functions, we examined embryos carrying the neomorphic mutation in the Arp1 subunit of the dynactin complex (*arp1^4D2^).* This mutant has been previously shown to affect the cortical anchoring of microtubules in the Drosophila oocyte without interfering with cargo transport (Nashchekin *et al.,* 2016). Additionally, the mutation results in an increase in both the growth and catastrophe rates of microtubule plus ends, resulting in shorter microtubules that fail to reach the posterior end of the Drosophila oocyte (Nieuwburg *et al*., 2017). To examine the requirement of the Dynactin complex specifically in the amnioserosa, we expressed Arp1 RNAi and Dynamitin in the amnioserosa using an amnioserosa specific Gal4.

### *The arp1^4D2^* mutation alters amnioserosa geometry and dynamics

We first examined the influence of the *arp1^4D2^* mutation on the contraction dynamics of the amnioserosa in embryos homozygous for a recombinant chromosome carrying the *arp1^4D2^*mutation and the Ubi::ECadherinGFP transgene which expresses ECadherin under the regulation of the ubiquitin promoter. Homozygous Ubi::ECadherinGFP embryos served as controls. Amnioserosa tissue contraction in both genotypes (assessed from 125 minutes prior to the completion of closure-t0, from maximum intensity projections of time-lapse movies) revealed that the area of the amnioserosa contained within the ellipse marked by ECadherin at the interface between the amnioserosa and the epidermis was smaller in mutant embryos as compared to controls (Fig. 1 A-B, G). Also, in contrast to control embryos in which the amnioserosa perimeter was shaped like an eye with canthi at its anterior and posterior ends (Fig. 1 A1-A7), the contour of the amnioserosa in the *arp1^4D2^* mutants was irregular (Fig.1 B2-B4), and the formation of the anterior canthus was delayed (Fig. 1 A1-A7, B1-B7, Movie S1). Consistent with this, the aspect ratio of the amnioserosa over the course of dorsal closure was also significantly increased in *arp1^4D2^* mutants compared to control embryos (Fig. 1H). This increase was primarily due to an increase in the width /intercanthal distance of the ellipse. These results reveal that the *arp1^4D2^*mutation alters the dynamics of amnioserosa geometry qualitatively and quantitatively. A prospective analysis of amnioserosa tissue areas from 12000 or 8000 μm^2^ to the end of closure however revealed no significant differences in constriction rates or dorsal closure duration between control and mutant embryos (Fig. S1 A, B).

**Figure 1.**
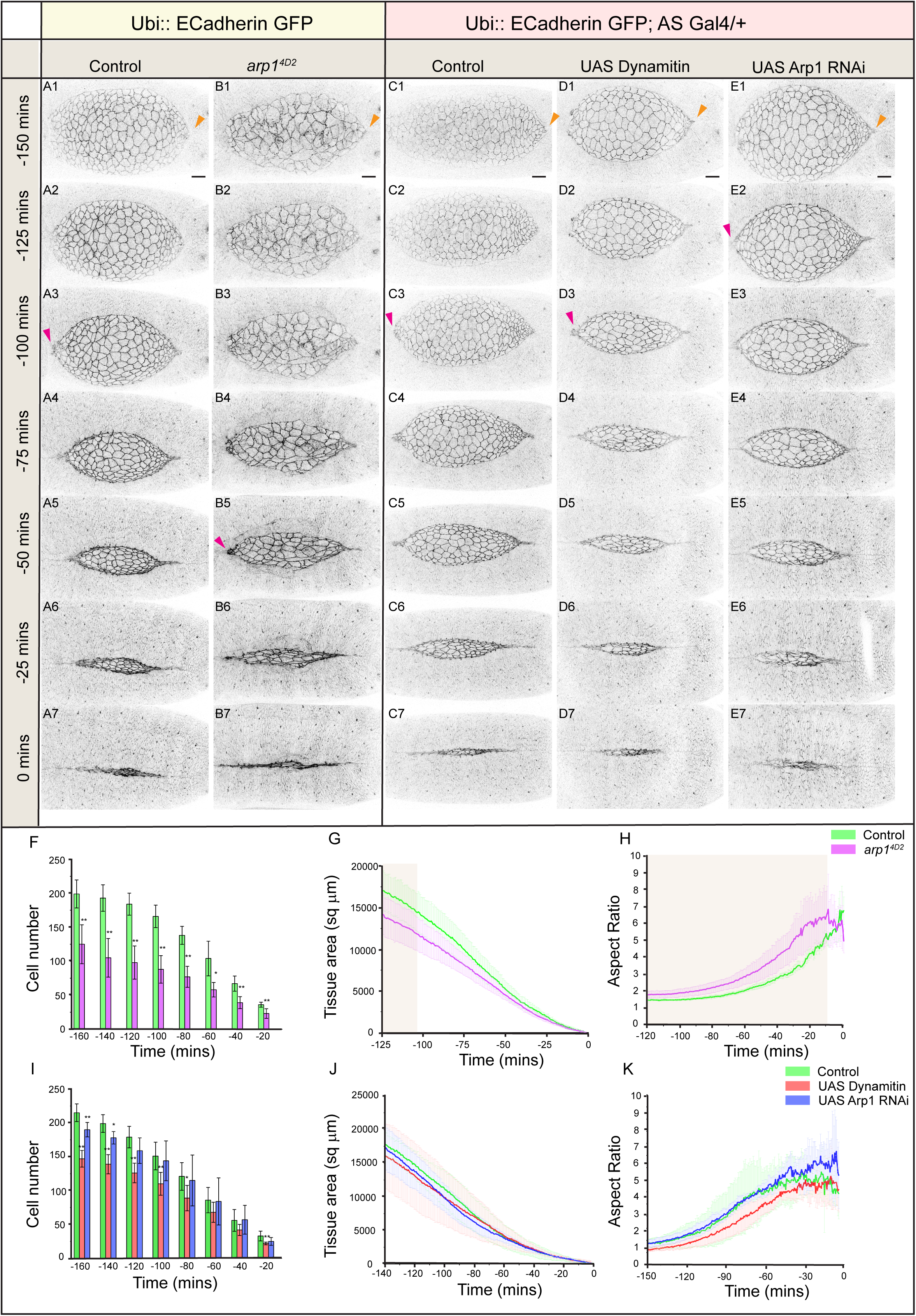
Dynactin perturbations influence tissue dynamics in the amnioserosa. (A-E) Snapshots of maximum intensity projections from real-time confocal microscopy images showing the progress of dorsal closure in embryos that are homozygous for the Ubi::ECadherin transgene (control, A1-A7) or are additionally homozygous for the *arp1^4D2^* mutation (B1-B7), are heterozygous for the Ubi::ECadherin transgene (control, C1-C7) or additionally express Dynamitin (D1-D7) or Arp1 RNAi (E1-E7) in the amnioserosa. 0 mins marks the end of dorsal closure. Anterior is to the left, and the pink and orange arrowheads point to the formation of the anterior and posterior canthi. (F, I) Average number of amnioserosa cells per embryo in the 120 minutes prior to the completion of closure in control (green, n=6) and *arp1^4D2^* mutant (magenta, n=6) embryos (F) or in control (green) embryos and in embryos expressing Arp1 RNAi (blue) or Dynamitin (red) in the amnioserosa (I). (G, J) Amnioserosa tissue area dynamics in the 120 minutes prior to the completion of closure in control (green, n=6) and *arp1^4D2^*mutant (magenta, n=6) embryos (G) or in control (green) embryos and in embryos expressing Arp1 RNAi (blue) or Dynamitin (red) in the amnioserosa (J). (H, K) Amnioserosa tissue aspect ratio dynamics in the 120 minutes prior to the completion of closure in control (green, n=6) and *arp1^4D2^* mutant (magenta, n=6) embryos (H) or in control (green) embryos and in embryos expressing Arp1 RNAi (blue, n=5) or Dynamitin (red, n=5) in the amnioserosa (K). Scale bar: 20 μm. See also Figure S1 and Movies S1 and S2.

To determine whether the morphodynamic effects observed in the *arp1^4D2^*mutants are due to the requirement of Arp1 function specifically in the amnioserosa, we downregulated Arp1 in the amnioserosa by expressing Arp1 RNAi using an amnioserosa-specific Gal4. We also expressed Dynamitin, which has been shown to irreversibly disrupt the Dynein-Dynactin complex (Burkhardt *et al*., 1997), in the amnisoerosa to determine whether Arp1 functions are dependent on the Dynein-Dynactin complex. In contrast to *arp1^4D2^* mutants, both the amnioserosa-specific perturbations tested exhibited no significant qualitative or quantitative defects in amnioserosa size, shape or dynamics including the formation of canthi (Fig. 1 C-E, J-K, Movie S2). These results suggest that the residual function of the complex due to maternal stores or insufficient downregulation, or the combination of a reduced amnioserosa area and the seemingly defective canthus formation might allow closure kinetics to proceed relatively unperturbed in the perturbations tested.

### Perturbations that affect Dynein-Dynactin complex function alter amnioserosa cell number

We surmised that the reduction in amnioserosa tissue area and altered amnioserosa geometry observed in *arp1^4D2^* mutants might result from defects in cell number, cell constriction or cell adhesion. We first examined the number of amnioserosa cells during the course of dorsal closure. In control embryos, amnioserosa cell numbers decrease approximately ten-fold from approximately 200 at 160 minutes prior to closure, to 20 at 20 minutes prior to closure. In contrast, the amnioserosa in *arp1^4D2^*mutants showed only a two-fold reduction in the number of cells, and a change in the temporal pattern and rate of decrease. Notably, amnioserosa cell numbers in the mutants were significantly lower than in controls (Fig.1 A-B, F, Movie S1). A similar but less severe reduction in amnioserosa cell numbers was also observed in Arp1 knockdown and Dynamitin overexpressing embryos particularly during early dorsal closure (Fig.1 I, Movie S2). These observations suggest that the Dynein-Dynactin complex might influence amnioserosa cell division prior to dorsal closure.

### Perturbations that affect Dynein-Dynactin complex function alter apical constriction dynamics in the amnioserosa

To determine whether amnioserosa cell constriction might also be influenced by the function of the Dynein-Dynactin complex, we segmented and tracked Ubi::ECadherin GFP-tagged amnioserosa cells (as described above) for 120 minutes prior to closure in both control and *arp1^4D2^* mutant embryos. In control embryos, most of the amnioserosa cells exhibit a polygonal (5-7 sides with a hexagonal predominance) cross section, whose apical surface area decreases over the period of dorsal closure (Pope and Harris, 2008). This area reduction happens in two phases. Amnioserosa cells in control embryos have larger areas and exhibit high amplitude oscillations with little or no net reduction in Phase I, and show reduced oscillation amplitudes and increased net constriction in Phase II (Fig. 2 A1-A5). In *arp1^4D2^* mutants, amnioserosa cells were larger in size than in control embryos, but their rate of constriction was significantly higher in both phases (Fig. 2 A-D, S1 A-B). Although the normalized pulse amplitude of amnioserosa cells (quantified for a duration of 30 minutes during pulsed apical constriction / Phase I; Solon *et al*., 2009; Blanchard *et al*., 2010; Saravanan *et al.,* 2013) in *arp1^4D2^* mutant embryos was not statistically different from that of the controls, amnioserosa cells in these mutants exhibited a wider range of pulse amplitudes, indicating a possible defect in the patterning of apical constriction or increased deformability (Fig. 2F). The latter possibility is consistent with the observation that amnioserosa cell shapes in *arp1^4D2^* mutant amnioserosa cells were highly irregular apically, with finger-like extensions that often project into the neighboring cells. This is in stark contrast to cells from control embryos whose apical interfaces become progressively straighter with time (Fig. 2 A1-A5, B1-B5, C1-C5, Movie S1). An increase in mean amnioserosa apical cell areas in phase I of dorsal closure (from 140 to 90 mins before closure) was also observed in the amnioserosa-specific knockdown of Arp1 and upon overexpression of Dynamitin both in real-time (Fig. 2 E) and in fixed preparations of stage-matched control and mutant embryos (Fig. S2 C-G). Surprisingly, unlike the mutants, both amnioserosa-specific perturbations demonstrated a significant reduction in the normalized pulsed amplitude (Fig. 2F). Whether the observed differences between the *arp1^4D2^*mutant and the amnioserosa-specific perturbations of the Dynein-Dynactin complex result from reduced loss of function, or whether they indicate different spatial requirements of the complex remains to be resolved. Nonetheless, these results point to the requirement of Arp1 and Dynactin complex functions in the modulation of cell deformability.

**Figure 2:**
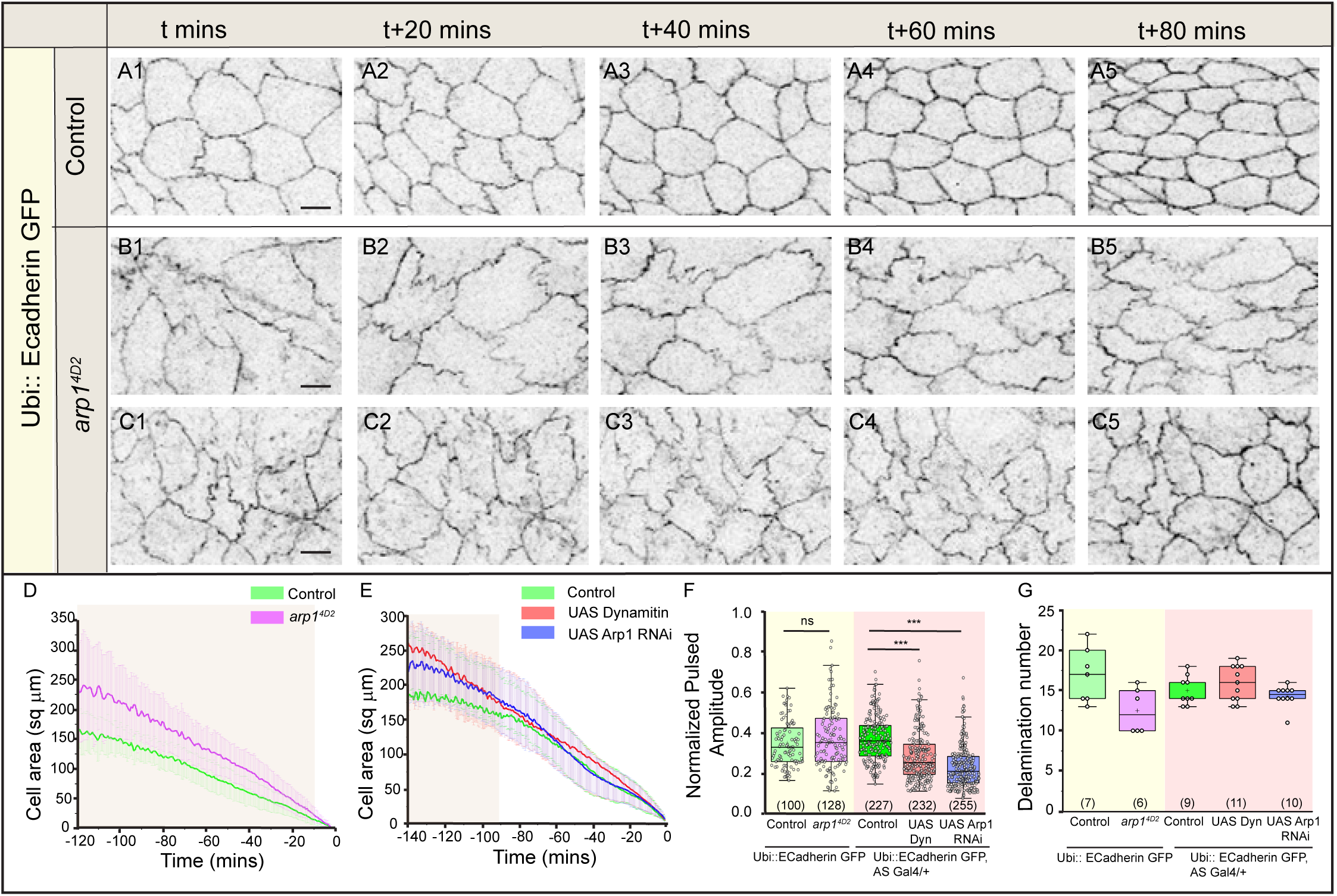
Dynactin perturbations influence amnioserosa cell morphodynamics during dorsal closure. (A-C) Snapshots of maximum intensity projections from real-time confocal microscopy images of showing amnioserosa cell morphodynamics in dorsal closure stage embryos that are homozygous for the Ubi::ECadherin transgene (controls, A1-A7) or are additionally homozygous for the *arp1^4D2^* mutation (B1-B7, C1-C7). (D,E) Cell area dynamics in the 120 minutes prior to the completion of closure in embryos that are homozygous for the Ubi::ECadherin transgene (D, control, green, n=34 cells from 3 embryos) or are additionally homozygous for the *arp1^4D2^*mutation (D, mutant, magenta, n=30 cells from three embryos) and in embryos that are heterozygous for the Ubi::ECadherin transgene (E, control, green, n=35 cells from four embryos) or additionally express Dynamitin (E, red, *n*=55 cells from five embryos) or Arp1 RNAi (E, blue, *n*=48 cells from five embryos) in the amnioserosa. Shaded areas in D and E indicate significant differences between control and mutants. (F) Normalised pulse amplitude in control amnioserosa cells and in the dynactin perturbations (numbers in brackets show number of cells analysed from at least 3 embryos). (G) Number of delaminations in control embryos and in embryos carrying the dynactin perturbations (numbers in brackets show number of embryos analysed). Mann-Whitney test and an unpaired Student’s *t*-test (***P<0.001, **P<0.01, *P<0.05) were used for statistical analysis. Scale bar: 10 μm. See also Fig. S2.

### The Dynein-Dynactin complex influences the morphodynamics of amnioserosa cell delamination

As described earlier, the amnioserosa is post-mitotic during the time course of dorsal closure, and there is no cell addition. Amnioserosa cells are, however, lost through cell delamination. In control embryos, delamination is a rare and seemingly stochastic cell behaviour in that its location and timing cannot be determined a priori (Kiehart *et al*., 2000; Toyama *et al*., 2008; Muliyil *et al.,* 2011). Earlier work from the lab uncovered the differential reorganization of the apical microtubule meshwork in the delaminating cell and its nearest neighbours. While apical microtubules appeared to be reduced in the delaminating cell, they were aligned and polarized in the nearest neighbours, with their plus ends towards the interfaces with the delaminating cell (Meghana *et al*., 2011; Guru *et al*., 2022). It seemed plausible that the microtubules in the neighbours may be anchored to the cortex underlying its interfaces with the delaminating cell, and that this may depend on the Dynein-Dynactin complex. We therefore examined delamination propensity in the *arp1^4D2^*mutants which, as described above, also showed a reduction in amnioserosa cell numbers. No statistically significant differences were observed in the amnioserosa cell delamination frequency between control embryos, the *arp1^4D2^*mutants and amnioserosa-specific dynactin perturbations examined (Fig. 2G). However, we observed differences in the dynamics of cell delamination.

The majority of the delaminating cells in control embryos constricted nearly isotropically (symmetric) to form a multi-way vertex (70%, n=118) that rarely resolved by junction growth (1%, n=118) during the course of dorsal closure, while a smaller fraction (29%, n=118) constricted anisotropically predominantly along one axis (asymmetric) to result in the formation of a new bicellular junction between two cells that were not neighbors before delamination (Fig. S3 A1-A5, F). In contrast, the percentage of symmetric delaminations that did not resolve was reduced in the *arp1^4D2^*mutants (27%, n=75), while percentage of asymmetric delamination events that resolved through the formation of a bicellular junction increased (53 %, n=75). Also, while the majority of multi-way vertices formed after symmetric delamination in controls were stable for at least 30 minutes (symmetric unresolved; Fig. S3 A1-A5, F), those in *arp1^4D2^* mutants resolved to form a *de-novo* junction (symmetric-resolved/ T2 transition) in 20% of delaminating cells (n=75) compared to a negligible number of such events (1 %, n=118) in controls (Fig. S3 B1-B5, F). Similar differences in the dynamics of delamination, with an increase in the fraction of both asymmetric delaminations and delaminations accompanied by resolution (25% in Dynamitin overexpression, n=170 and 31% in Arp1 RNAi, n= 143 compared to 4% in controls, n=135; Fig. S3 C-F) were observed in embryos expressing Arp1 RNAi or Dynamitin. Whether the observed phenotypes reflect endogenous differences in delamination dynamics between the genotypes, or whether they reflect phenotype modification in the presence of the Ubi::ECadherinGFP transgene remains to be determined. Nonetheless, these findings suggest that Dynein-Dynactin complex functions modulate adherens junction remodelling during amnioserosa cell delamination in sensitized genetic backgrounds.

### Altered ECadherin levels and distribution in *arp1^4D2^* mutants

Consistent with a role in adherens junction remodelling, the dynein heavy chain (DHC), a component of the Dynein-Dynactin complex was found to be enriched in the apical cortex of the amnioserosa cells at the level of the adherens junctions where it marked stippled apical cell outlines in control embryos. This enriched cortical localization of dynein heavy chain (DHC) colocalizes with ECadherin (Fig. 3 A1-A3). Intense spots of DHC spatially coincided with similarly increased intensities/heterogeneities in ECadherin. This apparent colocalization was better visualized in the line intensity plots, where most of the ECadherin peaks (representing the membrane, magenta) exhibited a corresponding DHC (green) peak (Fig. 3 C). Less intense puncta were found in the cytoplasm throughout the amnioserosa (Fig. 3 A2). Consistent with a function for Arp1 in the Dynein-Dynactin complex in amnioserosa cells, this cortical enrichment of DHC was lost in *arp1^4D2^* mutant embryos (Fig. 3 B1-B3, F). Additionally, the colocalization of the DHC and ECadherin peaks was also lost (Fig. 3D).

**Figure 3.**
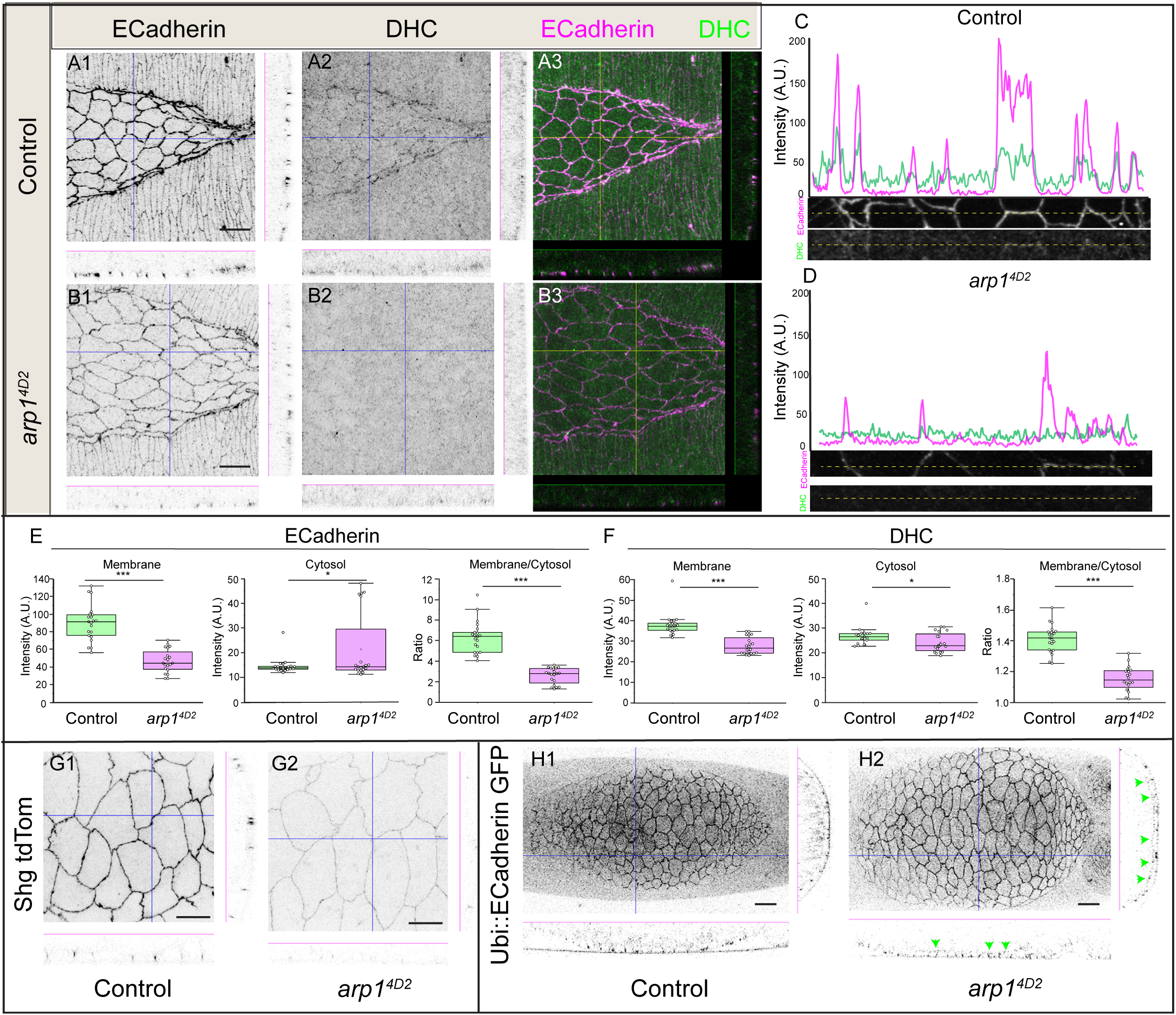
Dynactin perturbations influence ECadherin distribution. (A, B) Apical projections and orthogonal XZ and YZ slices of the amnioserosa during dorsal closure in control (A1-A3) and *arp1^4D2^* mutant (B1-B3) embryos, stained with ECadherin (magenta in A3, B3; grey in A1, B1) and DHC (green in A3, B3; grey in A2, B2). (C, D) Intensity profiles for ECadherin (magenta) and DHC (green) in control (C) and *arp1^4D2^* (D) embryos along the stippled line indicated in the image ROIs below it. (E, F) Box plots showing normalized intensity of ECadherin (E) and DHC (F) in the apical membrane and in the cytosol and their ratio measured from apical projections of the amnioserosa in control (n= 21 cells from five embryos) and *arp1^4D2^* mutant (n=20 cells from five embryos) embryos. (G1, G2) Maximum intensity apical projections and orthogonal XZ and YZ slices (made along the lines shown in the projection) of amnioserosa cells from embryos homozygous for the shg tdTomato transgene without (G1, n=8 embryos) or with (G2, n=11 embryos) the *arp1^4D2^* mutation. (H1, H2) Maximum intensity projections and orthogonal XZ and YZ slices (made along the lines shown in the projection) of embryos expressing homozygous ECadherinGFP without (H1, N=11 embryos) or with (H2, N=14 embryos) *arp1^4D2^*embryos (green arrows indicate basal ECadherin). In the box plots in E and F, boxes show median (horizontal line) ± interquartile range, the mean is indicated by ‘+’. Mann-Whitney test was used for statistical analysis. (*P<0.05, ***P<0.001). Scale bar: 10 μm (A, B, G) and 20μm (H).

The levels of ECadherin in the subapical plasma membrane were reduced in the *arp1^4D2^* mutants along with a marginal increase in the cytosolic pool (Fig. 3 A1, B1, C-E). ECadherin expression from both the Ubiquitin promoter driven Ecadherin GFP and the endogenous promoter driven Shg tdTomato transgenes was also reduced in *arp1^4D2^* mutants. A basal enrichment of ECadherin (marked with green arrowheads) in addition to the apical reduction was evident in the amnioserosa from the orthogonal sections in the former (Fig. 3 H). The differences in Ecadherin localization observed in the presence of transgenes producing different amounts of ECadherin (higher in Ubi::ECadherinGFP) suggest that Dynactin mutations may also be sensitive to ECadherin dosage.

### Tissue dynamics in *arp1^4D2^* mutants is sensitive to ECadherin levels

To understand the influence of altered ECadherin dosage, we examined amnioserosa tissue dynamics in *arp1^4D2^* mutant embryos carrying the Shg tdTomato transgene. A gross examination of tissue morphology of these mutants revealed more severe phenotypes compared to those carrying Ubi-Ecadherin GFP. While mutant embryos carrying Ubi::ECadherin GFP exhibited no significant effects on dorsal closure progression or outcome, other than a delay in anterior canthus formation, dorsal closure failed to complete in more than half of the mutant embryos carrying Shg tdTomato which also exhibited defects in amnioserosa integrity (53%, n=9/17, Fig. 4 A-E). In the majority of the mutant embryos that failed to complete closure, tears in the amnioserosa originated at sites of cell delamination (37%, n=6/17 embryos; Fig. 4 C, E), while the remainder originated elsewhere (16%, n=3/17 embryos; Fig. 4 D-E, Movies S3-S5). In the subset of embryos that successfully completed dorsal closure (47%, n=8/17, Movies S3), amnioserosa tissue area dynamics, inferred from the retrospective quantitative analysis of amnioserosa morphodynamics 140 minutes prior to closure, was indistinguishable from control embryos (Fig. 4F). These findings suggest a crucial role for the Dynein-Dynactin complex in the regulation of tissue integrity, possibly through the tuning of ECadherin levels during its contraction.

**Figure 4.**
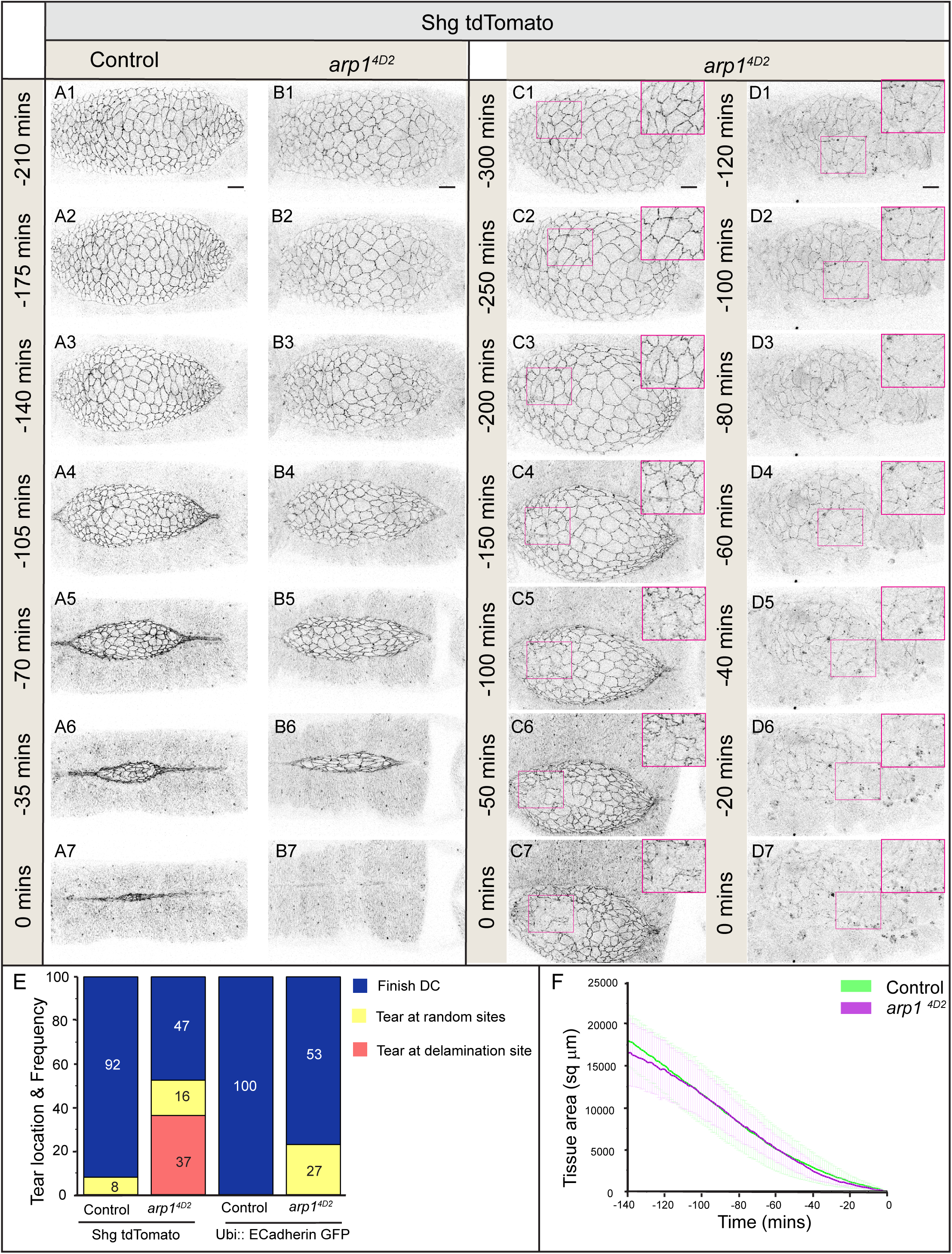
Tissue dynamics in dynactin perturbations is sensitive to ECadherin levels. (A-D) Snapshots of maximum intensity projections from real-time confocal microscopy images showing the progress of dorsal closure in embryos that are homozygous for the shg tdTomato transgene (control, A1-A7) or are additionally homozygous for the *arp1^4D2^* mutation (B-D). 0 mins marks the end of closure in A, B and the arrest of closure in C, D. 9 out of 17 *arp1^4D2^* mutant embryos fail to complete closure (C, D) due to amnioserosa tears at the site of delamination (inset in C) or elsewhere (inset in D). (E) Distribution of dorsal closure outcomes (completion or failure with amnioserosa tears, classified by site of origin) in control and *arp1^4D2^* mutant embryos carrying either Ubi::ECadh GFP or shg tdTom. (F) Amnioserosa tissue area dynamics in the 120 minutes prior to the completion of closure in control (green, n=6) and *arp1^4D2^* mutant (magenta, n=6) embryos carrying the shg tdTomato transgene. Mann-Whitney test was used for statistical analysis. Scale bar:20μm. See Movies S3, S4.

### Cell morphodynamics in *arp1^4D2^* mutants is sensitive to ECadherin levels

We further investigated the consequences of reduced ECadherin levels on cellular phenotypes associated with *arp1^4D2^* mutant embryos carrying the Shg tdTomato transgene. No significant differences in cell constriction rates or pulse amplitude were observed between control embryos and the subset of *arp1^4D2^*mutants that completed dorsal closure (Fig 5 F, G). Nonetheless, cells in these *arp1^4D2^*mutant embryos exhibited highly irregular shapes characterize by large lobules that protruded into neighbouring cells, and appeared qualitatively more deformable than their Ubi::ECadherin GFP expressing counterparts (Fig. 5 D-E, Movie S6). In the *arp1^4D2^* embryos carrying the Shg tdTomato transgene which displayed more severe tissue phenotypes, the delamination frequency was significantly reduced (Fig. 5H), and an increase in asymmetric delamination (43% in *arp1^4D2^* vs 13% in controls) was observed (Fig. 5I). Of the 57% of *arp1^4D^* mutant delaminations that were symmetric in *arp1^4D2^*(compared to 87% in control embryos), 10% exhibited resolution of the multi-way vertex while 2% failed to form a multi-way vertex and resulted in a small tear that disappeared as zippering progressed (Fig. 5I). These results suggest that adhesion remodelling that accompanies delamination is modulated by the Dynein-Dynactin function. Whether and how differential Dynein-Dynactin complex functions govern the coordinated regulation of cell adhesion in the delaminating cell and its nearest neighbours mechanisms remains to be discovered.

**Figure 5.**
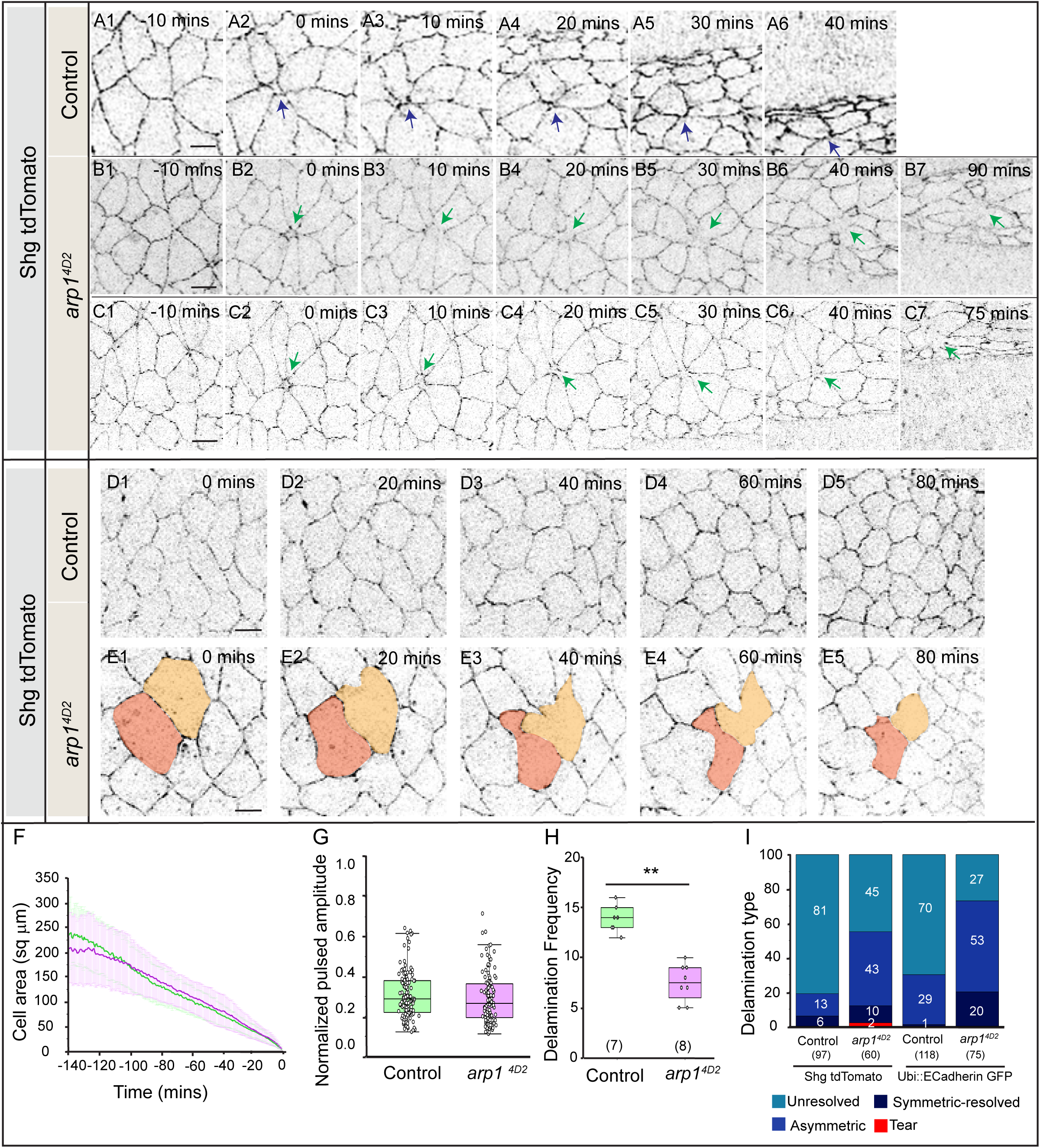
Cell morphodynamics in dynactin perturbations is sensitive to ECadherin levels. (A-C) Time-lapse images of maximum intensity projections showing morphodynamic features accompanying amnioserosa cell delamination in embryos expressing homozygous ShgtdTom without (A1-A6) or with (B1-B7, C1-C7) *arp1^4D2^*. Blue and green show delamination outcomes in control and mutant amnioserosa cells respectively. (D-E) Time-lapse images of maximum intensity projections showing morphodynamic features accompanying apical constriction in amnioserosa cells in embryos expressing homozygous ShgtdTom without (D1-D5) or with (E1-E5) the *arp1^4D2^* mutant. The coloured cells in E point to abnormal cell deformation. (F-H) Apical cell area dynamics (F), normalized pulse amplitude (G) and delamination frequency (H) of amnioserosa cells in control (light green, n=57 cells, in F, G and n=7 embryos in H) or *arp1^4D2^* mutant (magenta, n=44 cells, in F, G; n=8 in H) embryos carrying the ShgtdTom transgene. (I, J) Percentage distribution of delamination dynamics and outcomes (symmetry, resolution, tears) in control and *arp1^4D2^* mutant embryos carrying either Ubi::ECadh GFP (I) or shg tdTom (J). In the box plots in F and G, boxes show median (horizontal line) ± Interquartile range, the mean is indicated by ‘+’, and the sample size is in brackets. Mann-Whitney test (**P<0.01) was used for statistical analysis. Scale bar:10 μm. See also Fig. S3 and Movies S5, S6.

## Discussion

In this work, we discover roles for the Dynein-Dynactin complex in patterning cell morphodynamics accompanying epithelial fusion, a diversely deployed morphogenetic transformation whose failure compromises embryonic viability or produces significant disability. Using multiple genetic ways of perturbing the functions of this complex, we uncover the influence of this complex in the regulation of cell number, cell area, cell dynamics, and cell adhesion. Our findings allow us to suggest that the Dynactin complex functions to fine-tune and keep cell morphodynamic processes in check, at least in part through its effects on the regulation of cell adhesion. The mechanistic basis of its influence remains to be discovered.

One conspicuous effect observed in many of the genetic perturbations tested was the reduction in amnioserosa cell numbers, which was significantly different from controls only during early dorsal closure. This reduction could stem from the effects of this complex on either cell proliferation during early embryogenesis or on the maintenance of the differentiated state. The Dynein-Dynactin complex plays an essential role in cell division, and its localization at centrosomes has been shown to be essential not only for microtubule anchorage and spindle positioning but also for the spatiotemporally regulated enrichment of cell cycle regulators there (Skop and White, 1998; Quintyne and Schroer, 2002). While our work has not uncovered its basis, the apparent ‘recovery’ in cell numbers we observe in the amnioserosa specific perturbations later during dorsal closure suggests that feedback mechanisms, including, potentially, the reduction in cell delamination frequency, might reset the balance. Indeed, cell delamination is a cell behavior whose regulation maintains both cell number and tissue tension homeostasis (Marinari *et al*., 2012; Levayer *et al.,* 2016).

Dynactin complex perturbations also resulted in increased amnioserosa cell apical areas. This increase, like the decrease in cell numbers, was most evident during early closure in the amnioserosa specific perturbations. Whether this increase results from a failure of apical constriction or whether it reflects a compensatory response to reduced cell numbers, as has been documented elsewhere (Sonal *et al*., 2014), remains to be determined.

A striking phenotype associated with the amnioserosa cells, especially in the *arp1^4D2^* mutant, was their irregular shapes characterized by prominent lobular projections into the neighbours. This was in contrast to the early squiggly apical perimeter contours of wildtype amnioserosa cells that became progressively straighter. Cell deformability impacts the material properties of tissues and can have important implications for tissue dynamics (Saito and Ishihara, 2024). Uncovering the basis of the Dynactin complex dependent regulation of cell deformability, and specifically whether microtubule anchorage might underlie it, will be both important and interesting to determine.

Our results also suggest a role for the Dynactin complex in fine-tuning cell adhesion. Not only was the dynein heavy chain enriched in the same location as ECadherin, but perturbations in Dynactin complex function led to a reduction in ECadherin and were sensitive to ECadherin levels. While the reduction was evident throughout the amnioserosa, cell adhesion-dependent phenotypes were evident during cell delamination and in *arp1^4D2^* mutants and were modified by the ECadherin transgenes. Tears at delamination sites leading to compromised amnioserosa integrity and poor dorsal closure outcomes were seemingly ‘rescued’ by the ubiquitin promoter-driven ECadherin transgene, which provided higher than endogenous levels of ECadherin. In contrast, resolution of the multi-way vertex following delamination, rarely seen in control embryos, was facilitated by higher ECadherin levels. Pioneering work on cell intercalation accompanying germband extension in Drosophila has shed light on the nature and regulation of adherens junction remodelling accompanying junction shrinkage and junction growth and its interplay with the actomyosin cytoskeleton (Bertet *et al.,* 2004; Mira-Osuna and Borgne, 2024).

Our earlier work established the differential regulation of both the actomyosin and microtubule cytoskeleton in the delaminating cell and its nearest neighbours (Meghana *et al*., 2011). Whether and how the Dynactin complex is differentially regulated in the two cell populations and how it regulates adhesion remodelling will be interesting avenues to explore.

The effect of Dynactin complex perturbations on tissue dynamics was marginal. While this might be a consequence of residual function of the complex provided by maternal contribution or insufficient downregulation, the effect on tissue geometry hints at a requirement for ‘zipping’ at the canthus. Pioneering work has established the tissue and cellular contributions of forces that drive dorsal closure and the impacts of perturbing these forces on tissue geometry (Kiehart *et al*., 2000; Hutson *et al*., 2003; Solon *et al*., 2009). The Dynein heavy chain has also previously been shown to be enriched at the interface between the amnioserosa and the epidermis, especially close to the canthi (Eltsov *et al*., 2015). It will be interesting to determine whether and how the dynactin complex might also govern zipping and forces generated by amnioserosa contraction during dorsal closure.

## Conclusions

Our work uncovers the requirement for the Dynactin complex in modulating cell morphodynamics that accompany amnioserosa contraction during Dorsal closure whose mechanistic underpinnings remain to be discovered.

## Limitations of the study

While the phenotypic similarities observed upon perturbing the Dynactin complex in multiple ways suggests its requirement, stronger tissue specific perturbations and rescue, as well as other function modifying perturbations will be necessary to establish their direct or precise roles.

Our work identifies the Dynactin complex as a regulator cell morphodynamics and cell adhesion but does not resolve the mechanistic basis for its requirement. Additional genetic and molecular interactions studies, mosaic analysis, real-time imaging for quantitative morphometry and quantitative intensity measurements will enable a better delineation of the influence of the complex on cell deformability, cell adhesion remodelling and actin and microtubule cytoskeletal organisation.

## Acknowledgements

We thank Daniel St.Johnston, Tadashi Uemura, the Bloomington Drosophila Stock Centre (BDSC), the Vienna Drosophila Resource Centre (VDRC) and the Developmental Studies Hybridoma Bank (DSHB) for stocks and reagents, and members of the MN lab for discussion. We acknowledge support from TIFR/DAE, India (RTI4003 to MN and a graduate fellowship to DS).

## Author contributions

Deepanshu Sharma: Formal analysis; Investigation; Visualization; Methodology; Writing - original draft; Writing - review and editing; Performed all experiments, data generation and statistical analysis.

Maithreyi Narasimha: Conceptualization; Resources; Formal analysis; Supervision; Funding acquisition; Visualization; Methodology; Writing - original draft; Writing - review and editing; Conceived project, designed experiments and methodology, wrote first draft with DS, revised and edited manuscript, supervised the project and acquired funding

## Materials and Methods

### Materials

#### A. Reagents

| Primary Antibodies | Source | Identifier |
| --- | --- | --- |
| DCAD2<br>(ECadherin) | DSHB | AB_528120 |
| DHC | DSHB | AB_2091523 |
| Secondary Antibodies | Source | Identifier |
| Goat anti-Mouse IgG,<br>AlexaFluor 488 | Thermo Fisher Scientific | AB_2534088 |
| Goat anti-Rat IgG,<br>Cy5 | Thermo Fisher Scientific | AB_2534034 |
| Dyes & Drugs | Sources | Identifier |
| Vectashield Antifade<br>mounting medium | Vector Laboratories | Vector Laboratories_H- 1000 |

#### B. Drosophila stocks

| Experimental<br>Models:<br>Organisms/Strains | Sources | Identifier |
| --- | --- | --- |
| *D. melanogaster*:<br>Ubi::ECadherinGFP | (Uemura *et al.*, 1996)<br>Kind gift of Tadashi<br>Uemura. | NA |
| *D. melanogaster*:<br>UAS Arp1 RNAi | Bloomington Drosophila<br>Stock Centre (BDSC) | BDSC_67932 |
| *D. melanogaster*:<br>UAS DCTN2-p50 | Bloomington Drosophila<br>Stock Centre (BDSC) | BDSC_8784 |
| *D. melanogaster*:<br>Shg tdTom | Bloomington Drosophila<br>Stock Centre (BDSC) | BDSC_58789 |
| *D. melanogaster*:<br>*arp1*<sup>4D2</sup> | (Nieuwburg *et al.*, 2017)<br>Kind gift of Daniel St.<br>Johnston | NA |
| *D. melanogaster*:<br>332.3 Gal4 | Bloomington Drosophila<br>Stock Centre (BDSC) | BDSC_5398 |
| *D. melanogaster*:<br>c381 Gal4 | Bloomington Drosophila<br>Stock Centre (BDSC) | BDSC_3734 |
| *D. melanogaster*:<br>*w*<sup>1118</sup> | Bloomington Drosophila<br>Stock Centre (BDSC) | BDSC_5905 |
| UAS p150 Delta | Bloomington Drosophila Stock Center (BDSC) | BDSC_51645 |
| UAS p150 RNAi | Bloomington Drosophila Stock Center (BDSC) | BDSC_24760 |

#### C. Software and algorithms

| Software and Algorithms | Sources | Identifier |
| --- | --- | --- |
| FIJI | NIH | SCR_002285 |
| Tissue Analyzer (Fiji plugin) | (Aigouy *et al.*, 2016) | https://grr.gred-clermont.fr/labmirouse/software/WebPA/ |
| Adobe Photoshop | Adobe | SCR_014199 |
| Adobe Illustrator | Adobe | SCR_010279 |
| GraphPad | Prism | http://www.graphpad.com |
| OriginPro |  | RRID:SCR_014212 |

### Methods

#### A. Drosophila husbandry, stocks

The following mutant and transgenic lines obtained from the Bloomington Drosophila Stock Center: 332.3 Gal4 (for live imaging) and c381 Gal4 (for fixed preparations) for driving expression in the amnioserosa (AS Gal4); UAS Arp1 RNAi, UAS Dynamitin, UAS p150 RNAi, UAS p150 Delta for perturbing Dynactin complex functions and Shg tdTomato (to mark adherens junctions). Ubi::ECadhGFP was a kind gift of from Tadashi Uemura, Kyoto University, Japan (Uemura *et al*., 1996). *arp1^4D2^*was a kind gift from Daniel St Johnston, The Gurdon Institute, UK (Nieuwburg *et al*., 2017). The *arp1^4D2^* recombinants with Ubi::ECadherinGFP or Shgtd Tomato were built for this study. Crosses were designed to enable the unambiguous identification of the desired genotypes. The figure-wise list of genotypes analysed is provided below.

#### B. Genotypes List

**Figure 1**

The following genotypes were used to visualize and quantify amnioserosa tissue area, tissue aspect ratio and cell numbers in control embryos and in embryos carrying Dynactin perturbations.

w; Ubi::ECadherin GFP/ Ubi::ECadherin GFP

w; Ubi::ECadherin GFP/Ubi::ECadherin GFP; *arp1^4D2^*/*arp1^4D2^*

w; Ubi::ECadherin GFP, 332.3 Gal4/ +

w; Ubi::ECadherin GFP, 332.3 Gal4/ UAS Dynamitin.

w; Ubi::ECadherin GFP, 332.3 Gal4/ UAS Arp1 RNAi

**Figure 2**

The following genotypes were used to visualize and quantify apical amnioserosa cell area, pulsed apical constriction amplitude and constriction rate, as well as delamination frequency in control embryos and in embryos and in embryos carrying Dynactin perturbations.

w; Ubi::ECadherin GFP/ Ubi::ECadherin GFP

w; Ubi::ECadherin GFP/Ubi::ECadherin GFP; *arp1^4D2^*/*arp1^4D2^*

w; Ubi::ECadherin GFP, 332.3 Gal4/ +

w; Ubi::ECadherin GFP, 332.3 Gal4/ UAS Dynamitin.

w; Ubi::ECadherin GFP, 332.3 Gal4/ UAS Arp1 RNAi

**Figure 3**

The following genotypes were used in immunofluorescence and real-time experiments to visualise and quantify the levels and localization of Ecadherin and Dynein Heavy Chain.

For immunofluorescence

*w^1118^*

w;; *arp1^4D2^*/ *arp1^4D2^*

For real-time

w; shg tdTomato/shg tdTomato

w; shg tdTomato/shg tdTomato; *arp1^4D2^*/ *arp1^4D2^*

w; Ubi::ECadherin GFP/ Ubi::ECadherin GFP.

w; Ubi::ECadherin GFP/Ubi::ECadherin GFP; *arp1^4D2^*/*arp1^4D2^*

**Figure 4**

The above genotypes were used to visualize and quantify amnioserosa tissue area and tear location and frequency in control embryos and in embryos carrying *arp1^4D2^* mutant.

w; shg tdTom/shg tdTom

w; shg tdTom/shg tdTom; *arp1^4D2^*/ *arp1^4D2^*

w; Ubi::ECadherin GFP/ Ubi::ECadherin GFP

w; Ubi::ECadherin GFP/Ubi::ECadherin GFP; *arp1^4D2^*/*arp1^4D2^*

**Figure 5**

The above genotypes were used to visualize and quantify amnioserosa cell area, pulse amplitudes, delamination frequency and outcome, in control embryos and in embryos carrying the *arp1^4D2^* mutant.

w; shg tdTom/shg tdTom

w; shg tdTom/shg tdTom; *arp1^4D2^*/ *arp1^4D2^*

w; Ubi::ECadherin GFP/ Ubi::ECadherin GFP

w; Ubi::ECadherin GFP/Ubi::ECadherin GFP; *arp1^4D2^*/*arp1^4D2^*

**Figure S1**

The following genotypes were used to visualize and quantify apical amnioserosa tissue area in control embryos and in embryos carrying the *arp1^4D2^* mutant.

w; Ubi::ECadherin GFP/ Ubi::ECadherin GFP

w; Ubi::ECadherin GFP/Ubi::ECadherin GFP; *arp1^4D2^*/*arp1^4D2^*

**Figure S2**

The following genotypes were used for immunofluorescence to visualize and quantify amnioserosa apical cell area in control embryos and in embryos carrying Dynactin perturbations

*w^1118^*

w;; *arp1^4D2^*/ *arp1^4D2^*

w; UAS Arp1 RNAi/+;; c381 Gal4/+

w; UAS Dynamitin/+;; c381 Gal4/+

w; UAS p150 RNAi/+;; c381 Gal4/+

w; UAS p150 Delta/+;; c381 Gal4/+

**Figure S3**

The following genotypes were used to visualize and quantify delamination types and frequency in control embryos and in embryos carrying Dynactin perturbations.

w; Ubi::ECadherin GFP/ Ubi::ECadherin GFP

w; Ubi::ECadherin GFP/Ubi::ECadherin GFP; *arp1^4D2^*/*arp1^4D2^*

w; Ubi::ECadherin GFP, 332.3 Gal4/ +

w; Ubi::ECadherin GFP, 332.3 Gal4/ UAS Dynamitin.

w; Ubi::ECadherin GFP, 332.3 Gal4/ UAS Arp1 RNAi

#### C. Embryo harvesting and staging

For genetic perturbations and their controls, flies were allowed to lay for 4 hours, and embryos were aged at 29°C for 8 hours to enrich for stages of dorsal closure. For the analysis of cell and tissue dynamics, embryos were chosen that had completed germband retraction but had not yet initiated the formation of the anterior canthus, and embryos were imaged until the completion of closure. Tissue dynamics was analyzed in the 140 min before the completion of closure, while cell dynamics was analyzed from 140-120 minutes prior to completion of dorsal closure till its end. For the analysis of cell delamination, embryos were analyzed from the formation of the posterior canthus until the end of closure. Cell numbers were counted from 160 min before the completion of closure till its end.

#### D. Live imaging

For live imaging, embryos were dechorionated using 50% bleach for 2 minutes and washed with water. Embryos were then put on a 0.17 mm coverslip (Corning) on a thin film of Halocarbon oil 700 (Sigma) and imaged on inverted microscopes (Olympus FluoView1000, Olympus FluoView1200 and Olympus FluoView3000 confocal microscopes) on a 60X, 1.4NA objective with a digital zoom of 1X. Optical sections 0.4-1.1µm apart were acquired at 2 frames/minute. Maximum intensity projections were made using the Image5D plug-in on ImageJ and assembled as a time series on ImageJ. Images were prepared in Adobe Photoshop, and figures were assembled using Adobe Illustrator.

#### E. Immunofluorescence

For fixed preparations, embryos were harvested and stained using standard protocols (Narasimha and Brown, 2006). The following primary antibodies were used: anti-GFP (rabbit 1:1000, Invitrogen), anti-ECadherin (rat 1:10; DSHB) and anti-DHC (mouse 1:50, DSHB). Alexa Fluor-conjugated secondary antibodies (Invitrogen) were used at 1:200 dilutions. Embryos were then stored and mounted in Vectashield, and z stacks of confocal images with optical slices at 0.3 µm intervals were taken on (Olympus FluoView1000, Olympus FluoView1200 and Olympus FluoView3000) confocal microscopes using a 60×1.4 NA objective. Maximum intensity projections were made using the Image5D plug-in on ImageJ. For dual color imaging, sequential excitation with 488 nm and 561 nm lasers was used. Images were prepared in Adobe Photoshop, and figures were assembled on Adobe Illustrator (Adobe Systems).

#### F. Image manipulation/post-acquisition processing

No manipulations other than level adjustments to stretch the intensity histogram (when necessary) were applied for visualization purposes.

#### G. Intensity measurements

For fixed preps, ECadherin and DHC intensities were measured from maximum intensity projections of apical slices of the central amnioserosa cells during dorsal closure from control or *arp1^4D2^* embryos that had been stained with anti-ECadherin and anti-DHC antibodies. For membrane intensity measurements, a segmented line with 5-pixel width was used as ROI in ImageJ, while for cytosolic intensity, polygonal ROIs (3 pixels internal to the membrane ROI) were used. The intensity ratio was calculated using Microsoft Excel and plotted using Origin. For line intensity profiles, a line passing through multiple cells was drawn, and the ECadherin and DHC intensities were plotted using MS Excel.

#### H. Quantitative morphodynamics

Apical areas of individual cells in phase I of dorsal closure were measured in frames starting from −140 or −120 minutes prior to closure till its completion from maximum intensity projected, time-lapse images of Ubi::ECadherinGFP or ShgtdTomato. Central amnioserosa cells were segmented using Tissue Analyzer (Aigouy *et al.,* 2016), the selected ROIs were exported to Fiji (Schindelin *et al*., 2012), and the areas extracted. Cell area dynamics were analyzed by plotting the changes in the average apical area (±s.d.) as a function of time (Microsoft Excel). Similarly obtained cell area traces were also used to measure the amplitudes of pulses (Saravanan *et al.,* 2013). Normalized pulse amplitude was calculated as (Amax/A0) - (Amin/A0), where Amax and Amin represent, respectively, the maximum and minimum apical cell areas during a pulse, and A0 denotes the initial cell area. Pulse amplitude was calculated from pulses observed in cells in phase I over a 30-minute interval and averaged over multiple cells from multiple embryos of the same genotype to obtain the mean and standard deviation. The area of the amnioserosa was measured as the area of the ellipse from 140 minutes before the completion of closure to its end from maximum intensity projected, time-lapse images of Ubi::ECadherinGFP or ShgtdTomato. The areas were extracted and plotted as mentioned above. The total number of cells was marked and counted using Fiji from 140 minutes before the completion of closure to the end of closure. The number of cell delamination events occurring in the time window from the formation of the posterior canthus formation to the end of dorsal closure was marked and counted using Fiji.

#### I. Qualitative morphodynamics of cell deamination

Cell delamination events were classified into four types based on new junction formation or remodeling patterns. Delaminating cells that maintained a multi-way vertex for atleast 30 min were classified as unresolved. Those that initiated the formation of a new junction within 30 minutes of the formation of a multi-way vertex were scored as resolved. Delaminating cells that showed no multi-way vertex but constricted asymmetrically to form a new bicellular junction were classified as asymmetric. Delamination events that failed to form a stable junction or multipoint vertex but progressed to form a gap or tear were classified as tears.

#### J. Graphs and statistical analysis

No statistical method was employed to predetermine the sample size. The sample size (N) represents data from multiple experiments (immunofluorescence staining) and imaging sessions (live imaging), each containing one or many biological replicates (embryos or cells) depending on the type of analysis performed (detailed in the sections above), as it was not possible to analyze a statistically significant number of embryos from a single imaging session. No data was excluded from the analysis. All measurements (individual data points) are presented in most of the graphs provided.

Box plots were made using Origin Pro. The boxes show median ± interquartile range. Thick black lines show the median, the plus mark shows the mean, and the box limits indicate 25th and 75th percentiles. Whiskers extend to 1.5 times the interquartile range from the 25th and 75th percentiles. Individual data points are represented by dots. The sample size (n) analyzed for the individual genotypes is mentioned in the graph. All statistical analysis was carried out using GraphPad Prism 5 and Microsoft Excel software. See accompanying Figure-wise **Statistical table** in Supplementary Files for details.

| Figure | Populations compared | Sample size | Statistical test | Normal distribution | Variances | p-value |
| --- | --- | --- | --- | --- | --- | --- |
| 1F (T=-20) | Control<br>*arp1<sup>4D2</sup>* | N=6<br>N=6 | Mann<br>Whitney<br>test | Yes<br>Yes | Equal | 0.0077<br>(**) |
| 1F (T=-40) | Control<br>*arp1<sup>4D2</sup>* | N=6<br>N=6 | Mann<br>Whitney<br>test | No<br>Yes | Equal | 0.005 (**) |
| 1F (T=-60) | Control<br>*arp1<sup>4D2</sup>* | N=6<br>N=6 | Mann<br>Whitney<br>test | Yes<br>Yes | Equal | 0.01 (*) |
| 1F (T=-80) | Control<br>*arp1<sup>4D2</sup>* | N=6<br>N=6 | Mann<br>Whitney<br>test | Yes<br>No | Equal | 0.0022<br>(**) |
| 1F (T=-100) | Control<br>*arp1<sup>4D2</sup>* | N=6<br>N=6 | Mann<br>Whitney<br>test | Yes<br>Yes | Equal | 0.0022<br>(**) |
| 1F (T=-120) | Control<br>*arp1<sup>4D2</sup>* | N=6<br>N=6 | Mann<br>Whitney<br>test | Yes<br>Yes | Equal | 0.005 (**) |
| 1F (T=-140) | Control<br>*arp1<sup>4D2</sup>* | N=6<br>N=6 | Mann<br>Whitney<br>test | Yes<br>Yes | Equal | 0.0022<br>(**) |
| 1F (T=-160) | Control<br>*arp1<sup>4D2</sup>* | N=6<br>N=6 | Mann<br>Whitney<br>test | Yes<br>Yes | Equal | 0.0022<br>(**) |
| 1I (T=-20) | Control<br>UAS Arp<br>RNAi | N=6<br>N=6 | Mann<br>Whitney<br>test | Yes<br>Yes | Equal | 0.08 |
| 1I (T=-40) | Control<br>UAS Arp<br>RNAi | N=6<br>N=6 | Mann<br>Whitney<br>test | Yes<br>Yes | Equal | 0.9271 |
| 1I (T=-60) | Control<br>UAS Arp<br>RNAi | N=6<br>N=6 | Mann<br>Whitney<br>test | Yes<br>Yes | Equal | 0.5368 |
| 1I (T=-80) | Control<br>UAS Arp<br>RNAi | N=6<br>N=6 | Mann<br>Whitney<br>test | Yes<br>Yes | Equal | 0.6623 |
| 1I (T=-100) | Control<br>UAS Arp<br>RNAi | N=6<br>N=6 | Mann<br>Whitney<br>test | Yes<br>Yes | Equal | 0.8548 |
| 1I (T=-120) | Control<br>UAS Arp<br>RNAi | N=6<br>N=6 | Mann<br>Whitney<br>test | Yes<br>Yes | Equal | 0.0814 |
| 1I (T=-140) |  |  |  |  | Equal |  |
|  | Control<br>UAS Arp<br>RNAi | N=6<br>N=6 | Mann<br>Whitney<br>test | Yes<br>Yes |  | 0.0353<br>(*) |
| 1I (T=-160) | Control<br>UAS Arp<br>RNAi | N=6<br>N=6 | Mann<br>Whitney<br>test | Yes<br>Yes | Equal | 0.0087<br>(**) |
| 1I (T=-20) | Control<br>UAS Dyn | N=6<br>N=6 | Mann<br>Whitney<br>test | Yes<br>Yes | Equal | 0.0035<br>(**) |
| 1I (T=-40) | Control<br>UAS Dyn | N=6<br>N=6 | Mann<br>Whitney<br>test | Yes<br>No | Equal | 0.08 |
| 1I (T=-60) | Control<br>UAS Dyn | N=6<br>N=6 | Mann<br>Whitney<br>test | Yes<br>Yes | Equal | 0.132 |
| 1I (T=-80) | Control<br>UAS Dyn | N=6<br>N=6 | Mann<br>Whitney<br>test | Yes<br>Yes | Equal | 0.026 (*) |
| 1I (T=-100) | Control<br>UAS Dyn | N=6<br>N=6 | Mann<br>Whitney<br>test | Yes<br>Yes | Equal | 0.0087<br>(**) |
| 1I (T=-120) | Control<br>UAS Dyn | N=6<br>N=6 | Mann<br>Whitney<br>test | Yes<br>Yes | Equal | 0.0049<br>(**) |
| 1I (T=-140) | Control<br>UAS Dyn | N=6<br>N=6 | Mann<br>Whitney<br>test | Yes<br>Yes | Equal | 0.005 (**) |
| 1I (T=-160) | Control<br>UAS Dyn | N=6<br>N=6 | Mann<br>Whitney<br>test | Yes<br>Yes | Unequal | 0.0022<br>(**) |
| 1G (T-125 to T-105) | Control<br>*arp1<sup>4D2</sup>* | N=4<br>N=3 | Student'<br>s t test | Yes<br>Yes | Equal | <0.05(*) |
| 1H (T-120 to T-12) | Control<br>*arp1<sup>4D2</sup>* | N=4<br>N=3 | Student'<br>s t test | Yes<br>Yes | Equal | <0.05(*) |
| 1J (T-140 to T=0) | Control<br>UAS Dyn | N=4<br>N=5 | Student'<br>s t test | Yes<br>Yes | Equal | >0.05 |
| 1J (T-140 to T=0) | Control<br>UAS Arp<br>RNAi | N=4<br>N=5 | Student'<br>s t test | Yes<br>Yes | Equal | >0.05 |
| 1K (T-150 to T=0) | Control<br>UAS Dyn | N=4<br>N=5 | Student'<br>s t test | Yes<br>Yes | Equal | >0.05 |
| 1K (T-150 to T=0) | Control<br>UAS Arp<br>RNAi | N=4<br>N=5 | Student'<br>s t test | Yes<br>Yes | Equal | >0.05 |
| 2D (T-120 to T-6) | Control<br>*arp1<sup>4D2</sup>* | N=4,<br>n=34 | Student'<br>s t test | Yes<br>Yes | Equal | <0.05(*) |
|  |  | N=3,<br>n=30 |  |  |  |  |
| 2E (T-140 to T-90) | Control<br>UAS Dyn | N=4,<br>n=35<br>N=5,<br>n=55 | Student'<br>s t test | Yes<br>Yes | Equal | <0.05(*) |
| 2E (T-140 to T-90) | Control<br>UAS Arp<br>RNAi | N=4,<br>n=35<br>N=5,<br>n=48 | Student'<br>s t test | Yes<br>Yes | Equal | <0.05(*) |
| 2F | Control<br>*arp1<sup>4D2</sup>* | N=4,<br>n=100<br>N=3,<br>n=128 | Mann<br>Whitney<br>test | No<br>Yes | Equal | 0.3092 |
| 2F | Control<br>UAS Dyn | N=4,<br>n=227<br>N=5,<br>n=232 | Mann<br>Whitney<br>test | Yes<br>No | Equal | 8.689121<br>E<sup>-15</sup> (***) |
| 2F | Control<br>UAS Arp<br>RNAi | N=4,<br>n=227<br>N=5,<br>n=255 | Mann<br>Whitney<br>test | Yes<br>Yes | Equal | 1.4065E<sup>-37</sup> (***) |
| 2G | Control<br>*arp1<sup>4D2</sup>* | N=7<br>N=6 | Mann<br>Whitney<br>test | Yes<br>Yes | Equal | 0.0726 |
| 2G | Control<br>UAS Dyn | N=9<br>N=11 | Mann<br>Whitney<br>test | Yes<br>Yes | Equal | 0.3537 |
| 2G | Control<br>UAS Arp<br>RNAi | N=9<br>N=10 | Mann<br>Whitney<br>test | Yes<br>No | Equal | 0.7057 |
| 3E (membrane) | Control<br>*arp1<sup>4D2</sup>* | N=5,<br>n=21<br>N=5,<br>n=20 | Student'<br>s t test | Yes<br>Yes | Equal | 9.263E<sup>-10</sup> (***) |
| 3E (cytosol) | Control<br>*arp1<sup>4D2</sup>* | N=5,<br>n=21<br>N=5,<br>n=20 | Student'<br>s t test | No<br>No | Equal | .0353(*) |
| 3E(membrane/cyto<br>sol) | Control<br>*arp1<sup>4D2</sup>* | N=5,<br>n=21<br>N=5,<br>n=20 | Student'<br>s t test | Yes<br>No | Unequal | 1.192654<br>E<sup>-10</sup> (***) |
| 3F (membrane) | Control<br>*arp1<sup>4D2</sup>* | N=5,<br>n=21<br>N=5,<br>n=20 | Student'<br>s t test | No<br>Yes | Equal | 5.356E<sup>-8</sup> (***) |
| 3F (cytosol) | Control<br>*arp1<sup>4D2</sup>* | N=5,<br>n=21<br>N=5,<br>n=20 | Student'<br>s t test | No<br>No | Equal | .0323(*) |
| 3F(membrane/cyto<br>sol) | Control<br>*arp1<sup>4D2</sup>* | N=5,<br>n=21<br>N=5,<br>n=20 | Student'<br>s t test | Yes<br>Yes | Equal | 1.341984<br>E<sup>-11</sup>(***) |
| 4F (T-140 to T0) | Control<br>*arp1<sup>4D2</sup>* | N=6<br>N=6 | Student'<br>s t test | Yes<br>Yes | Equal | >0.05 |
| 5F (T-140 to T0) | Control<br>*arp1<sup>4D2</sup>* | N=7,<br>n=57<br>N=8,<br>n=44 | Student'<br>s t test | Yes<br>Yes | Equal | >0.05 |
| 5G | Control<br>*arp1<sup>4D2</sup>* | N=7,<br>n=57<br>N=8,<br>n=44 | Student'<br>s t test | Yes<br>Yes | Equal | >0.05 |
| 5H | Control<br>*arp1<sup>4D2</sup>* | N=7<br>N=8 | Mann<br>Whitney<br>test | Yes<br>Yes | Equal | 0.0014<br>(**) |
| S1A | Control<br>*arp1<sup>4D2</sup>* | N=6<br>N=6 | Student'<br>s t test | Yes<br>Yes | Equal | >0.05 |
| S1B | Control<br>*arp1<sup>4D2</sup>* | N=6<br>N=6 | Student'<br>s t test | Yes<br>Yes | Equal | >0.05 |
| S2G | Control<br>*arp14D2*<br>UAS *Arp1*<br>RNAi<br>UAS<br>Dyn<br>UAS p150<br>RNAi<br>UAS p150<br>Delta | n=90;<br>N=7<br>n=99;<br>N=8<br>n=48;<br>N=6<br>n=69;<br>N=6<br>n=75;<br>N=6<br>n=58;<br>N=6 | Student'<br>s t test | Yes<br>Yes<br>Yes<br>Yes<br>Yes<br>Yes | Equal<br>Equal<br>Equal<br>Equal<br>Equal<br>Equal | <0.001<br><0.001<br><0.01<br><0.001<br><0.001 |

## Legends to Movies

**Movie S1:** Amnioserosa tissue dynamics accompanying dorsal closure in homozygous Ubi::ECadhGFP embryos with or without the *arp1^4D2^*mutation.

**Movie S2:** Amnioserosa tissue dynamics accompanying dorsal closure in heterozygous Ubi::ECadhGFP embryos that also additionally express Arp1 RNAi or Dynamitin in the amnioserosa.

**Movie S3:** Amnioserosa tissue dynamics accompanying dorsal closure in homozygous shgtdTomato embryos with or without the *arp1^4D2^*mutation

**Movie S4:** Amnioserosa tissue dynamics accompanying dorsal closure in homozygous shg tdTomato embryos with or without the *arp1^4D2^*mutation showing tears in the amnioserosa in the mutant.

**Movie S5:** Amnioserosa cell morphodynamics accompanying dorsal closure in homozygous shg tdTomato embryos with or without the *arp1^4D2^* mutation showing tears at the site of delamination in the mutant).

**Movie S6:** Amnioserosa cell delamination morphodynamics accompanying dorsal closure in homozygous shg tdTomato embryos with or without the *arp1^4D2^* mutation showing cell deformability in the mutant).

---

**Figure S1:**
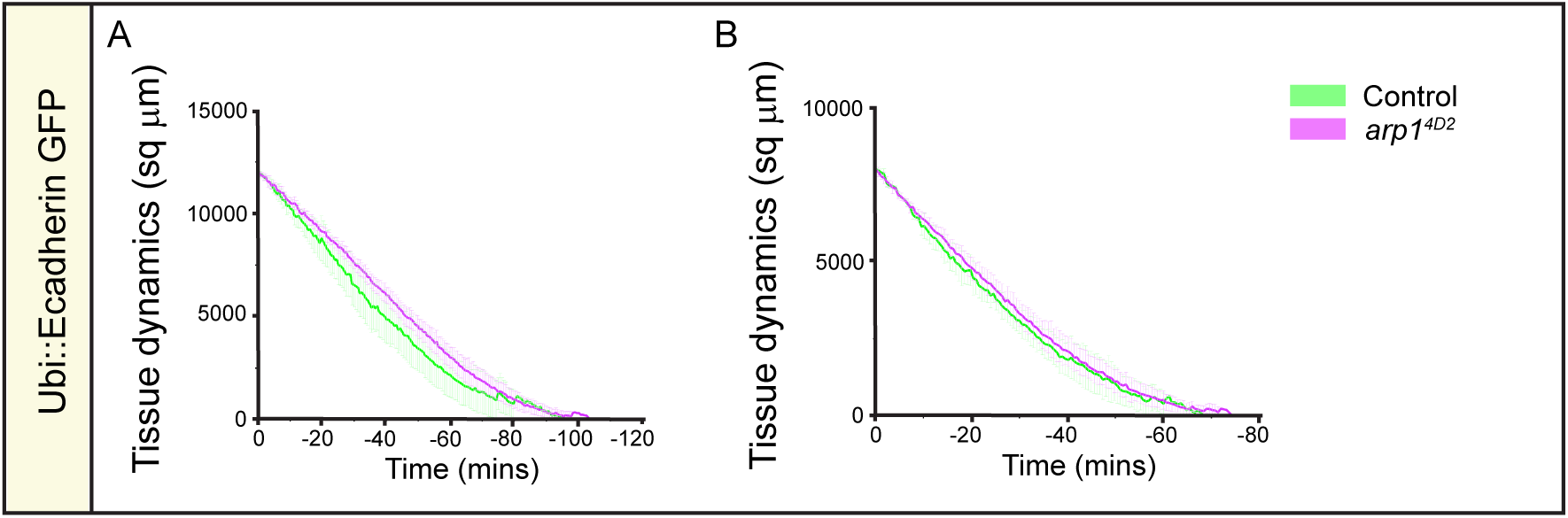
The duration of dorsal closure is not significantly affected upon perturbing dynactin. (A, B) Amnioserosa tissue area dynamics analysed prospectively from starting amnioserosa areas of 12000 (A) or 8000 (B) μm^2^ to ∼ 0 μm^2^ in embryos that are just homozygous for Ubi::ECadherin GFP (green in A, B; n=6) or are additionally homozygous for the *arp1^4D2^* mutation (magenta, n=6).

**Figure S2:**
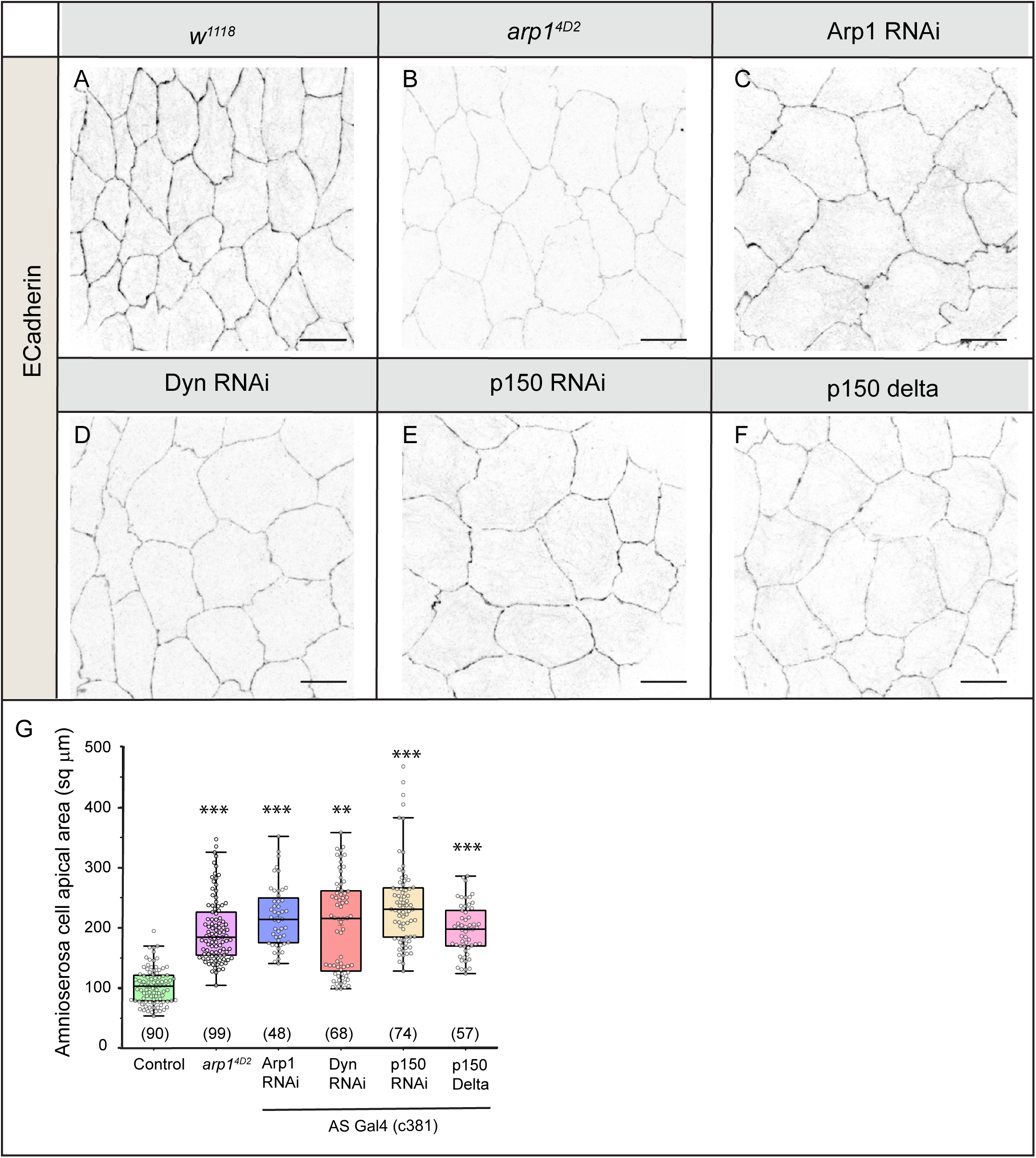
Dynactin perturbations influence amnioserosa cell area during dorsal closure. (A-G) Amnioserosa apical cell areas visualised (A-F) and quantified (G) in fixed preparations of stage-matched control embryos (A), homozygous *arp1^4D2^* mutants (B), and in embryos expressing Arp1 RNAi (C), Dynamitin (D), p150 RNAi (E) or p150 Delta (F) in the amnioserosa using AS Gal4 (c381). In the box plots in G, boxes show median +/− interquartile range, the mean is indicated by + and the sample size is in brackets. Mann-Whitney test was used for statistical analysis. ** P<0.01 and *** P<0.001. Scale bar: 10 μm.

**Figure S3:**
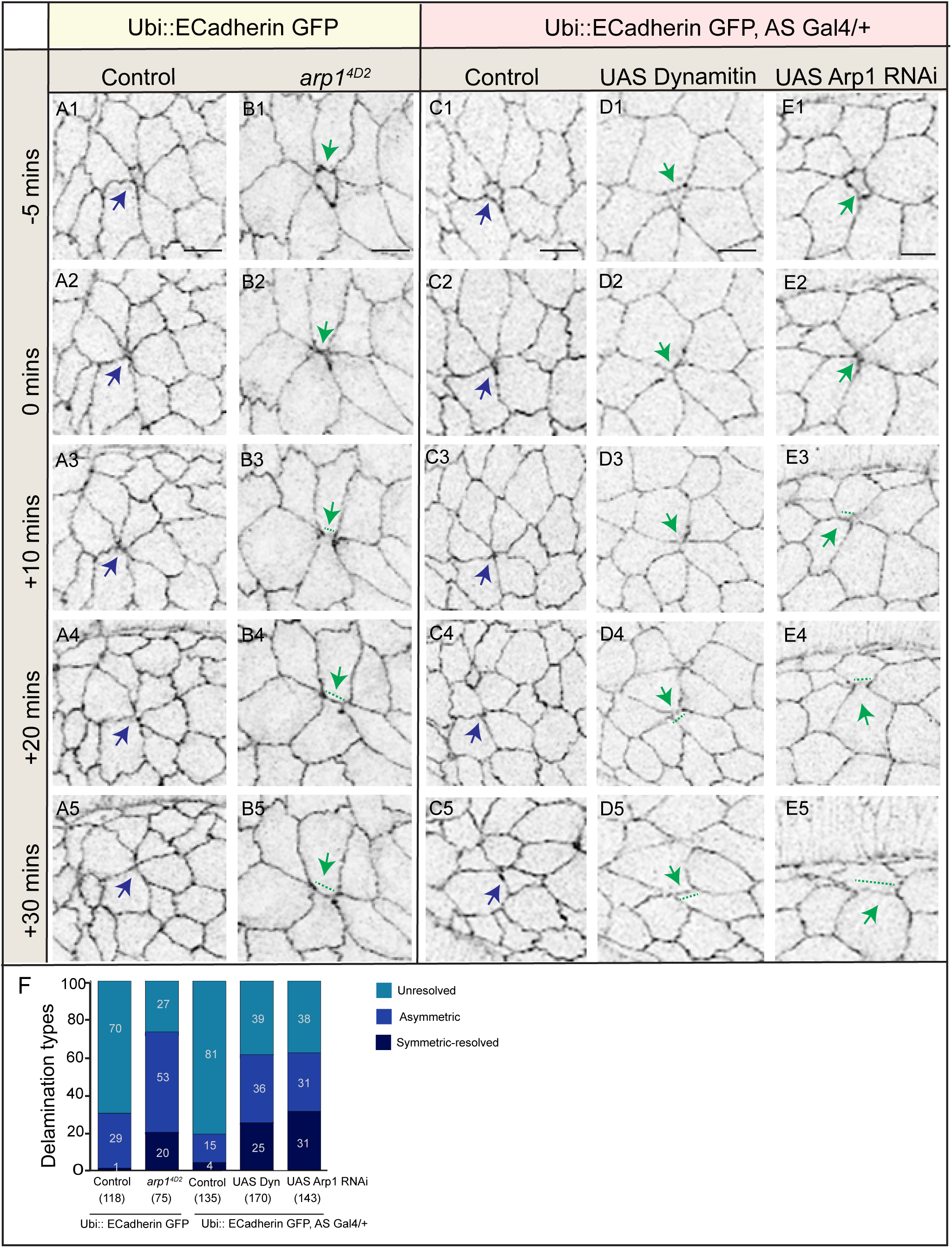
Dynactin perturbations influence amnioserosa cell delamination morphodynamics during dorsal closure. Snapshots of maximum intensity apical projections from real-time confocal microscopy images showing amnioserosa cell delamination morphodynamics in dorsal closure stage embryos that are homozygous for the Ubi::ECadherin transgene (controls, A1-A5) or are additionally homozygous for the *arp1^4D2^* mutation (B1-B5) and in embryos that are heterozygous Ubi::ECadherin transgene (C1-C5) and additionally express Dynamitin (D1-D5) or Arp1 RNAi (E1-E5) in the amnioserosa using AS Gal4 (332.3). t0 mins marks the point at which the apical area of the delaminating cell (blue, green arrowheads) is close to 0 μm^2^. (E) Percentage distribution of delamination dynamics and outcomes (symmetry, resolution, tears) in the above genotypes (numbers in brackets show number of delaminations analysed from at least 6 embryos). Scale bar: 10 μm.

